# Theta-paced stimulation of the thalamic nucleus reuniens entrains mPFC-HPC oscillations and facilitates the acquisition of extinction memories

**DOI:** 10.1101/2025.08.15.670624

**Authors:** Tuğçe Tuna, Flávio Afonso Gonçalves Mourão, Stephen Maren

**Affiliations:** Department of Psychological and Brain Sciences, Texas A&M University, College Station, TX; Institute for Neuroscience, Texas A&M University, College Station, TX; Beckman Institute for Advanced Science and Technology, University of Illinois Urbana-Champaign, Urbana, IL; Department of Psychology, University of Illinois Urbana-Champaign, Champaign, IL

## Abstract

**Background:** The nucleus reuniens (RE) is a midline thalamic nucleus interconnecting the medial prefrontal cortex (mPFC) and the hippocampus (HPC), structures known to be involved in aversive memory processes. Recent work indicates that the RE plays a critical role in the acquisition and retrieval of fear extinction memories. Functional inactivation of the RE impairs both mPFC-HPC coherence and extinction memory. Here we examine whether imposing theta activity on the RE entrains oscillations in the mPFC and HPC and facilitates extinction learning.

**Methods:** To deliver theta-paced (8 Hz) optogenetic stimulation, we expressed an excitatory opsin, (channelrhodopsin; AAV9-CaMKIIa-hChR2(H134R)-mCherry) or control virus (AAV9-CaMKIIa-mCherry) in the RE in male and female rats. A single optic fiber targeting the RE was implanted during the same surgery. After recovery, animals underwent auditory fear conditioning, extinction training, and an extinction retrieval test, each separated by 24-h. During extinction training, conditioned stimuli (CS) presentations were paired with 8-Hz sinusoidal optostimulation (473 nM, 10 mW) of the RE. In another experiment, we recorded local field potentials (LFPs) from the mPFC and dorsal HPC during using these behavioral procedures.

**Results:** Theta-paced stimulation of the RE during extinction training significantly decreased freezing behavior compared to the control group. Notably, the reduction in conditioned behavior was evident during the subsequent stimulation-free retrieval test. This reveals that RE stimulation during extinction not only suppresses conditioned fear responses acutely, but also facilitates the acquisition of long-term extinction memories. Theta-paced RE stimulation markedly enhanced both neural activity and entrained theta oscillations in the mPFC and dHPC.

**Conclusion:** This work suggests that the RE oscillatory activity is critical for the acquisition of extinction memories through the modulation of hippocampal-prefrontal network dynamics. In the future, RE theta-paced stimulation can be an important therapeutic tool by strengthening extinction memories.

## INTRODUCTION

Prolonged exposure-based therapies are commonly used to treat people with trauma- and stressor-related disorders, including post-traumatic stress disorder (PTSD). These therapies aim to extinguish fear by exposing people to the cues that trigger their trauma in a controlled therapy context [1, 2]. Extinction learning and memory, the underlying mechanism of exposure-based therapies, involve a complex circuitry including the medial prefrontal cortex (mPFC) and the hippocampus (HPC) [3–5].

Growing evidence has suggested a dynamic interplay between the mPFC and HPC, in which the HPC conveys contextual information to the mPFC, and the mPFC exerts top-down control over HPC-dependent memory retrieval processes [6–9]. Notably, while the HPC sends dense efferent projections to the mPFC, there are few direct return pathways from the mPFC to the HPC [10]. This asymmetry highlights the need for a hub for coordinating interactions between the mPFC and HPC. Accumulating evidence suggests that the nucleus reuniens (RE) is a potential hub for coordinating communication between the mPFC and HPC [6, 10–16]. The RE is a midline thalamic structure interconnecting the mPFC and HPC via its bidirectional projections with both structures [15, 17, 18]. The RE plays a critical role in encoding and retrieval of fear extinction memory [16, 19–25]. This role relies on its ability to modulate contextual memory processing by the regulation of synaptic plasticity in the HPC [26] and facilitating coordinated activity and synchronism between the mPFC and HPC [19, 21].

The RE synchronizes mPFC and HPC theta oscillations during extinction retrieval, and pharmacological inactivation of the RE impairs both extinction retrieval and the mPFC-HPC theta coherence. Furthermore, theta-paced (8-Hz) optogenetic stimulation of the RE prevents fear renewal following extinction, a common form of fear relapse [21]. However, it remains unknown if theta-paced RE stimulation entrains the mPFC and HPC network at theta oscillatory frequencies and if it can facilitate the acquisition of extinction memories.

In this study, we hypothesized that theta-paced RE stimulation would increase mPFC-HPC synchrony and facilitate the acquisition of extinction memories. To this end, we performed an auditory fear conditioning task followed by fear extinction training and an extinction retrieval test. Optogenetic stimulation of the RE at 8-Hz was applied during extinction training in adult male and female Long-Evans rats, specifically during the presentation of the conditioned stimuli (CS). Our results showed that RE theta-paced stimulation significantly reduced conditioned freezing behavior during extinction training. This effect was also evident during the stimulation-free retrieval test conducted the following day, suggesting that theta-paced stimulation of the RE suppresses conditioned freezing and facilitates the acquisition of long-term extinction memories. Moreover, in a subset of animals, mPFC and HPC local field potential (LFP) recordings were performed prior to conditioning and during extinction training, along with RE 8-Hz optogenetic stimulation. These recordings revealed a marked increase in oscillatory activity around 8-Hz in both the mPFC and dHPC, along with a significant increase in temporal coherence between these two structures. These findings suggest that oscillatory activity in the RE is essential to the acquisition of extinction memories by the coordination of hippocampal-prefrontal network dynamics.

## MATERIALS AND METHODS

### Animals

Adult male (*n* = 14) and female (*n* = 18) Long-Evans Blue Spruce rats (200-224g) purchased from Envigo (Indianapolis, IN, USA) were used for the experiments. Animals were acclimated to the vivarium for one day upon arrival and handled 1 min/day for at least five days before the experiments. They were single housed and kept at 14/10 h light/dark cycle with *ad libitum* access to food and water. All experiments were conducted during the light cycle and all procedures were approved by the Texas A&M University Animal Care and Use Committee and University of Illinois Urbana-Champaign Animal Care and Use Committee.

### Viruses

AAV9-CaMKIIa-hChR2(H134R)-mCherry and AAV9-CaMKIIa-mCherry viruses purchased from Addgene (Watertown, MA) were used for the optogenetics and electrophysiological recording experiments. Both viruses were diluted to a final titer of 5 × 10^12^ GC/mL with a Dulbecco’s phosphate buffered saline (DPBS) solution.

### Surgical procedures

Animals were anesthetized with isoflurane (5% for induction, ~2% for maintenance) and received i.p. injections of 5 mg/kg carprofen before being placed into a stereotaxic frame (Kopf Instruments; Tujunga, CA). The scalp was shaved and injected with lidocaine HCI (2%) with epinephrine (intradermal). Povidone-iodine was applied to the skin and an incision was made. The skull was leveled using the bregma and lambda coordinates.

For the optogenetics experiment, a hole was drilled above the nucleus reuniens (RE) (AP: −2.1 mm, ML: 1.25 mm, DV: −7.09 mm relative to the bregma surface with a 10° angle from the midline). The virus (0.5 μL) was infused at a rate of 0.1 μL/min using an injector connected to a Hamilton syringe mounted in an infusion pump (Kd Scientific; Holliston, MA) with polyethylene tubing. The injector tip was kept in place for an additional 10 min before being withdrawn. The incision was cleaned, sutured, and a topical antibiotic (Triple Antibiotic Plus, G&W Laboratories) was applied. Two weeks later, animals underwent a second surgery for fiber optic placement. Four small holes were drilled in the skull to affix four jeweler’s screws. The hole above the RE was reopened, and an optic fiber (10 mm, 200 μM core, 0.39 NA; RWD, China) was implanted (AP: −2.1 mm, ML: 1.25 mm, DV: −6.79 mm relative to the bregma surface with a 10° angle from the midline). The fiber was affixed to the skull with black dental cement (Contemporary Ortho-Jet Powder, Lang Dental).

For the electrophysiological recording experiment, animals underwent a surgery for virus injection as explained above. Two weeks later, they underwent the second surgery for fiber optic and multielectrode array placements (16-channel microwire arrays from Innovative Neurophysiology; Durham, NC). Four small holes were drilled in the skull to affix four jeweler’s screws. One of the screws was positioned posterior to lambda and functioned as a ground and reference. A stainless-steel wire was coiled and fixed around this screw with conductive silver paint (Silver Print II; GC Electronics). Three other holes were drilled, one above the RE (AP: −2.1 mm, ML: −1.25 mm, DV: −6.79 mm relative to the bregma surface with a 10° angle from the midline) to insert the fiber optic (10 mm, 200 μM core, 0.39 NA; RWD, China), and two others to insert the multielectrode arrays, above the mPFC (AP: 3.72 mm, ML: 0.75 mm, DV: −6.23 mm relative to bregma surface with a 15° angle) and the CA1 region of the dHPC (AP: −4.44 mm, ML: 3 mm, DV: −2.85 mm relative to bregma surface). The mPFC received an implant consisting of 16 tungsten wires arranged in a 2×8 matrix, with one row containing eight 8 mm electrodes and the other containing eight 6.9 mm electrodes. This design helped target simultaneously both the infralimbic (IL) and prelimbic (PL) cortices of the mPFC. The dHPC was implanted with a 4×4 array of 5-mm electrodes. All wires were spaced 200 μm apart, and each wire had a diameter of 50 μm. Multielectrode arrays and the fiber optic were affixed to the skull with black dental cement.

At the end of the surgeries, antibiotic ointment (Triple Antibiotic Plus, G&W Laboratories) was immediately applied to the incision to prevent bacterial infections. After recovery from anesthesia, animals were returned to the vivarium and allowed to recover for 7-10 days before beginning any behavioral and recording procedure.

### Behavioral procedures

For both experiments, animals underwent auditory fear conditioning, conditioning context exposure, extinction training, and extinction retrieval test (Fig. 1B) in conditioning chambers (30 x 24 x 21 cm, Med Associates, St. Albans, VT) in two different contexts. Context A was used for conditioning and conditioning context exposure, and Context B was used for extinction training and extinction retrieval testing. Context A involved transporting animals in white transport boxes without bedding. There were red room lights with cage lights, grid floors, and a black and white striped background in this context. In addition, 1.0 % ammonium was used to wipe the chambers. Context B involved black transport boxes with bedding, white room lights with no cage lights, plastic floors, and a black-and-white star-shaped background. 3.0 % acetic acid was used to wipe the chamber. Conditioning took place in Context A with five tone (CS: 10 s, 80 dB, 2 kHz) and foot shock (US: 1 s, 1.0 mA) pairings separated by 60 s interstimulus intervals (ISIs). To extinguish fear associated with the conditioning context, rats were exposed to Context A for 35 min the next day, in the absence of CS or US presentations. They then underwent cued extinction training in Context B with 45 CS-alone presentations separated by 30 s ISIs. 8-Hz theta-paced stimulation of the RE took place during extinction CSs. Blue light was delivered in an 8-Hz oscillating pattern using a waveform generator (RIGOL Technologies, Inc.; Portland, OR), which was driven by transistor-transistor logic (TTL) triggers from Med Associates software. Blue laser turned on five s before each CS and ended five s after, resulting in stimulation periods of 20 s in total. A stimulation-free extinction retrieval test took place the next day in Context B, with five CS-alone presentations separated by 30 s ISIs. For the electrophysiological recording experiment, animals first underwent a baseline recording session from the mPFC and dHPC in Context B with 8-Hz stimulations of the RE. Blue laser turned on after a three min laser-free baseline period and lasted 10 s. A total of five simulations were delivered separated by 30 s ISIs. All behavioral testing was separated by 24-h. A three-min baseline period was provided at the beginning of each testing session along with a post-trial period at the end of each session.

**Figure 1.**
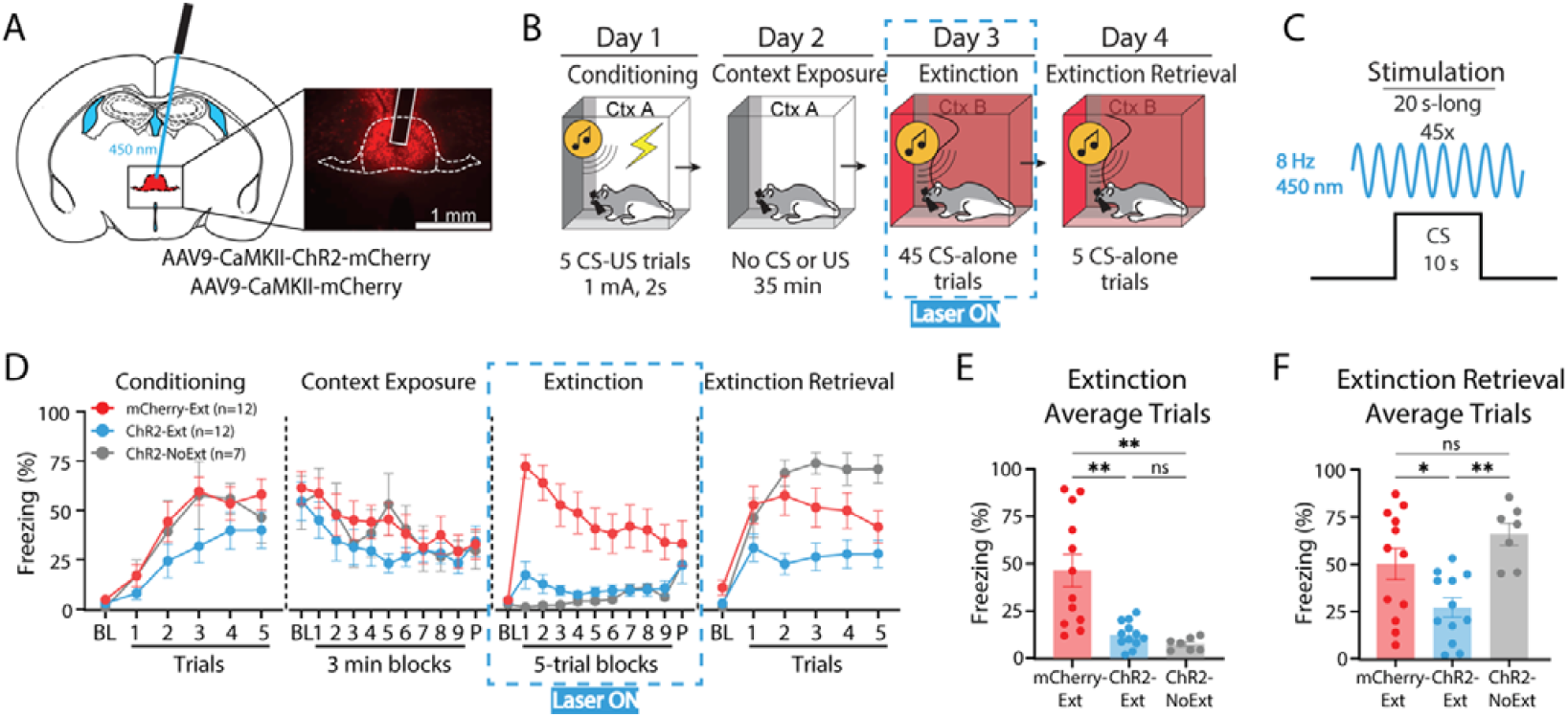
Optogenetic theta-paced stimulation of the RE facilitates extinction learning. **(A)** Surgery protocol (left) and a representative viral expression and fiber placement in the RE (right). AAVs encoding either the ChR2 or mCherry were injected and a single fiber optic was implanted in the RE approximately three weeks before the behavioral protocol. The RE was stimulated with a blue laser (450 nm). **(B)** Behavioral protocol used in the experiment. **(C)** Blue laser stimulation protocol used during extinction CSs. **(D)** Percent freezing levels averaged for groups in each of the behavior days. **(E)** Freezing levels averaged for extinction trials. With theta-paced stimulation, ChR2-Ext group (*n* = 12) showed significantly less freezing compared to mCherry-Ext group (*n* = 12) and did not differ from ChR2-NoExt group (*n* = 7) (Brown-Forsythe ANOVA: *F*_2, 12.5_ = 16.20, *p* =.0003, Dunnett’s T3 multiple comparison tests: mCherry-Ext vs ChR2-Ext: *p* =.007; mCherry vs ChR2-NoExt: *p* =.0029; ChR2-Ext vs ChR2-NoExt: *p* >.05). **(F)** Freezing levels averaged for extinction retrieval trials. ChR2-Ext group demonstrated facilitated extinction retrieval following theta-paced stimulation of the RE during extinction compared to both control groups (One-way ANOVA: *F*_2, 28_ = 7.268, *p* =.0029, Tukey’s multiple comparison tests: mCherry-Ext vs ChR2-Ext: *p* =.0443; ChR2-Ext vs ChR2-NoExt: *p* =.0029; mCherry-Ext vs ChR2-NoExt: *p* >.05). Data are represented as mean ± SEM.

For the optogenetics experiment animals and a subset of electrophysiological recording experiment animals (*n* = 2), motor activity was measured using a load-cell platform (Med Associates Inc.) and recorded with Threshold software (Med Associates). Freezing behavior was used as the index of fear and was defined as movement that is below 10% of the maximum movement signal for at least one second. For the remaining electrophysiological experiment animals (*n* = 6), freezing levels were assessed using a video-based recording system and analyzed with Video Freeze® software (Med Associates). In the Video Freeze® system, cameras were individually calibrated before each experimental animal, and freezing behavior was automatically quantified according to the default setting recommended by the manufacturer.

Using these camera recordings, we manually scored grooming behavior.

### Optogenetics

8-Hz stimulation was delivered with blue (450 nm) laser (Dragon Lasers). The laser power was calibrated to 10 mW at the optic fiber tip. 8-Hz sinusoidal stimulation protocol was created using the DG812 Waveform Generator (Rigol Technologies, Inc.; Portland, OR). TTL adapter triggers from the Med Associates software were used to control the laser.

### Electrophysiology recordings

Extracellular local field potential (LFP) activity was recorded using a multichannel electrophysiology system (Plexon; Dallas, TX). Briefly, from a monopolar design referenced to a skull screw, with other channels referenced to the LFP wire, the recorded wideband signal was amplified at 2000x and digitized at 40-kHz sampling rate.

### Histology

Animals were overdosed with sodium pentobarbital (Fatal Plus, 100 mg/mL, 0.5 ml, i.p) and those implanted with electrode arrays were given DC current lesions (0.1 mA pulse, 10s) (World Precision Instruments; Sarasota, FL) to mark electrode tips at the corners of the arrays. All animals were then transcardially perfused with ice-cold physiological saline followed by 10% formalin solutions. Brains were obtained, kept in 10% formalin solution at 4° C overnight and then transferred to a 30% sucrose solution until they sank (~3-4 days). 30μ-thick sections were obtained at −20° C using a cryostat (Leica Microsystems; Wetzlar, Germany). For the optogenetics experiment, sections were mounted and coverslipped using Fluoromont-G, with DAPI (Invitrogen; Carlsbad, CA) mounting medium. Viral expression and fiber optic placements were verified using a Zeiss microscope (Axio Imager). For the electrophysiology experiment, 30μ-thick sections were mounted on gelatine-subbed slides, stained with thionin (0.25%), and coverslipped using Permount (Fisher Scientific; Pittsburgh, PA) mounting medium. Electrode placements were verified using a brightfield microscope (Leica MZFLIII). Animals with missed injection, electrode, or fiber placements were excluded from further analyses.

### Grooming behavior

Grooming behavior was manually scored by two experienced raters who were blind to the experimental group of animals as well as the times in which 8-Hz stimulation took place. A custom-made software was used to mark the start and end timestamps for each grooming episode. The software was created in MATLAB^⍰^ 2024a and is publicly available in (https://github.com/marenlab). As with other reports [27–29], the start of each grooming episode was marked when rats started to perform elliptical strokes around their noses with front paws. The end of each grooming episode was marked when rats stopped grooming for at least 1 s as well as when they started to perform another behavior (e.g. freezing). We examined grooming behavior during stimulation-free baseline period, stimulation periods, and ISIs, for both baseline and extinction recording days, in a subset of electrophysiological recording animals (*n* = 6). Grooming duration percentage was calculated by dividing the time spent grooming in each of these phases by the total duration of these phases.

### LFP analysis

All analyses were conducted using a combination of built-in functions and custom code in MATLAB (https://github.com/marenlab). LFPs from a single channel were processed by detrending to ensure a constant mean and subsequently downsampled to 1 kHz. Segments containing motion artifacts and/or line noise were excluded from all analyses.

#### Power spectral density estimate

Power spectra were calculated using the pwelch.m function (Signal Processing Toolbox™) and set as 2s time-point, Hamming window, nfft = 2^15 and 90% overlapping. The data were normalized to the total power between 2-12 Hz and the average power around 8 (+/−0.2) Hz was then calculated to obtain a power estimate.

#### Wavelet spectrum

The illustrative time-frequency spectrum was calculated by a continuous 1-D wavelet transform on the *Z*-scored signal using cwt.m function (Signal Processing Toolbox™) set as the analytic Morse wavelet with ⍰θ = 60. The absolute values of the wavelet spectrum were normalized using *Z*-scores across the frequency domain to highlight relative changes in power over time, rather than merely reflecting absolute power differences between frequency bands.

#### Phase Coherence

The signals were first band-pass filtered between 7.5 and 8.5 Hz using the eegfilt.m function (EEGLAB; https://sccn.ucsd.edu/eeglab/). Phase angles were then extracted using the hilbert.m function (Signal Processing Toolbox™). Phase differences (Δ phase) were computed in 250 ms time windows by calculating the difference between the imaginary components of the respective brain regions.

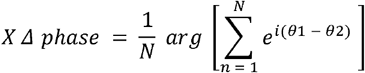

According to the equation, *X* Δ phase represents the argument of the sum of the phase difference vectors; *N* is the number of time samples for each signal; and *θ*□ and *θ*□ denote the phase values of the respective brain regions recorded. The phase-locking value (PLV) is a metric of phase coherence expressed as a dimensionless real number ranging from 0 to 1. It is calculated as the magnitude of the mean vector obtained from the distribution of phase differences. A PLV of 0 reflects a random or uniform phase distribution, while a PLV of 1 indicates perfect phase synchrony between the signals [30].

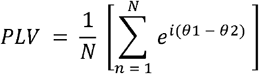

The data were represented as polar plots illustrating the distribution of phase differences and the magnitude of the mean vector (PLV) derived from this distribution. The illustrative PLV time-course was computed across the entire experimental session using 2-second sliding windows with a 95% overlap.

### Statistics

Behavioral data were represented as mean ± SEM. Repeated-measures analysis of variance (ANOVA) was followed by Tukey’s post hoc multiple comparisons test. Behavioral data from optogenetics and electrophysiological recording experiments were collapsed as the same behavioral protocol was used. Male and female data were collapsed where significant sex differences were not found. Prior work and literature determined the group sizes used in these experiments [19, 21]. *p*□<□0.05 was considered statistically significant. GraphPad Prism 9.5.0 was used for behavioral data analyses.

Electrophysiological data were expressed as median and interquartile range (25th–75th percentile). To evaluate the main effects, a non-parametric Aligned Rank Transform (ART) procedure was employed. For multiple comparisons, Wilcoxon signed-rank tests were used for within-group paired comparisons, and Mann–Whitney *U* tests (ranksum) were used for between-group comparisons. *p*□<□0.05 was considered statistically significant. Rosenthal’s correlation coefficient was used to estimate the effect size of statistically significant results. To further confirm the statistical significance, a permutation test was applied to both within-group and between-condition comparisons. A null distribution was generated by randomly shuffling insertion across 200 segments, repeated 1000 times. The observed values were then transformed into *Z*-scores relative to the null distribution, and the mean difference was calculated. All non-parametric statistical analyses were performed using custom code in MATLAB^⍰^ 2024a (https://github.com/marenlab).

## RESULTS

### Theta-paced stimulation of the RE facilitates extinction learning

The engagement of neural networks in oscillatory patterns can be influenced by the intrinsic resonance characteristics of individual neurons, which enhance their responsiveness to external rhythmic inputs at particular frequencies [31–33]. Accordingly, it has been demonstrated that optogenetic theta-paced stimulation of the RE at 8-Hz creates 8-Hz power in the RE and blocks the renewal of fear memory after extinction [21]. Here, we sought to answer if theta-paced stimulation of the RE also facilitates extinction learning. To this end, we injected adult male and female rats (*n* = 32; 14 male, 18 female) with adeno-associated viruses (AAVs) expressing either the blue light-shifted excitatory opsin Channelrhodopsin (ChR2) or a control fluorescent protein (mCherry) into the RE. The RE was also implanted with a single fiber optic (Fig. 1A. Also see Fig. S1 for viral expression and fiber optic placements).

After three weeks, animals underwent the fear conditioning protocol (Fig. 1B). All groups were conditioned with five tone (CS) and footshock (US) pairings in Context A. The next day, they were merely exposed to the conditioning context without CS or US presentations to extinguish the fear of the conditioning context. The next day, animals expressing the ChR2 virus (ChR2-Ext, *n* = 12, 5 males and 7 females) and those expressing the mCherry virus (mCherry-Ext, *n* = 12, 5 males and 7 females) were extinguished with 45 CS-alone trials in Context B. During each CS, 8-Hz blue light stimulation was delivered to the RE for 20 s-long periods, starting five seconds before the CS onset and ending five seconds after its offset (Fig. 1C). A third group of ChR2-expressing animals (ChR2-NoExt, *n* = 8) was exposed to the extinction context and received 8-Hz blue light stimulations, similar to those delivered in the other groups. However, this group was not exposed to the CSs and served as a no-extinction control, allowing us to isolate the effect of optogenetic stimulation from extinction itself. One animal from this no-extinction group was excluded from further analyses due to off-target viral expression (*n* = 7, 4 males and 3 females). Finally, all animals were tested for extinction retrieval with five CS-alone trials without RE stimulations.

During conditioning, the data for one squad of animals (*n* = 7) was inadvertently not recorded. However, all groups showed similar changes in freezing behavior over the course of the conditioning session (main effect of Trials: *F*_5, 105_ = 28.33, *p* <.0001) with no main effect of Group or Group x Trials interaction (*p*s >.05) (Fig. 1D, Conditioning). There was also no effect of sex or Sex x Trials interaction (*p*s >.05). The next day, during conditioning context exposure, all groups showed similar reductions in freezing, indicating fear of the conditioning context was extinguished (Fig. 1D, Context Exposure, main effect of Time blocks: *F*_10, 280_ = 5.105, *p* <.0001; no main effect of Group or Group x Trials interaction: *p*s >.05). Female rats exhibited lower freezing compared to male rats (main effect of Sex: *F*_1, 29_ = 5.157, *p* =.0308), but similar rates of learning (no Sex x Time blocks interaction: *F*_10, 290_ = 1.445, *p* =.1598). During extinction, however, group differences were observed with 8-Hz stimulation of the RE (Fig. 1D, Extinction, Group x Trials interaction: *F*_20, 280_ = 5.443, *p* <.0001). The ChR2-Ext group demonstrated significantly lower levels of freezing compared to the mCherry-Ext group (Tukey’s multiple comparison test, *p* =.0005). Moreover, freezing levels of the ChR2-Ext group were comparable to the ChR2-NoExt group, which was not presented with CSs in this novel context (Context B) (*p* =.8522). The mCherry-Ext group, on the other hand, showed significantly higher freezing compared to the ChR2-NoExt group (*p* =.0006). This was confirmed when the extinction trials were averaged (Fig. 1E). There were no sex differences in freezing during extinction (*p*s >.05).

Because a recent study indicated that optogenetic stimulation of the RE leads to compulsive-like grooming behavior [27], we examined grooming behavior in a subset of animals (*n* = 6, 2 ChR2, 4 mCherry). We performed 8-Hz stimulations during both a pre-conditioning baseline session and extinction CSs. As reported by [27], RE stimulation led to increased grooming behavior in ChR2 animals compared to mCherry animals during both baseline (Table S1) and early trials of extinction (Table S2, also see supplementary videos). Grooming behavior started shortly after the laser was turned on and usually exceeded the stimulation period and continued during ISIs. However, grooming behavior was not related to fear state. Animals showed similar grooming levels during both pre-conditioning baseline stimulation (low fear) and stimulation during the early trials of extinction (high fear). We also observed that in the later trials of extinction, one ChR2 animal no longer showed photostimulation-locked grooming while the other continued to groom (Table S3). The former rat showed improved extinction retrieval the next day while the latter exhibited higher freezing levels during extinction retrieval. Despite the small sample size, these results suggest that grooming behavior caused by RE photostimulation does not account for the effects of photostimulation on freezing to the CS during extinction or on the subsequent facilitation of extinction retrieval; these may be mediated by independent mechanisms.

During the stimulation-free extinction retrieval test, the ChR2-Ext group showed lower freezing levels compared to the other groups (Fig. 1D, Extinction Retrieval, Group x Trials interaction: *F*_10, 140_ = 4.979, *p* <.0001, Tukey’s post-hoc: ChR2-Ext vs ChR2-NoExt: *p* =.0069; ChR2-Ext vs mCherry-Ext: *p* =.0467), with no main effect of Sex or Sex x Trials interaction (*p*s >.05). Facilitation of extinction retrieval with RE 8-Hz stimulation was confirmed with averaged retrieval trials across group (*F*_2, 28_ = 7.268, *p* =.0029): ChR2-Ext group exhibited significantly less freezing compared to the mCherry-Ext (*p* =.0443) and ChR2-NoExt (*p* =.0029) groups, while the freezing level of mCherry-Ext group did not differ from the ChR2-NoExt group (*p* =.3109), which showed high freezing levels with CS presentations in Context B (Fig. 1F).

Overall, these results indicate that the theta-paced stimulation of the RE during extinction CSs suppresses conditioned freezing. The suppression in the conditioned freezing carries over to the stimulation-free retrieval test day, suggesting that the theta-paced stimulation of the RE facilitates the acquisition of longer lasting extinction memory. Our results suggest that theta-paced stimulation of the RE could ultimately be an important therapeutic tool by strengthening extinction memories.

### RE stimulation increases mPFC–dHPC theta oscillations

Theta oscillations in the mPFC and dHPC are engaged during the retrieval of fear extinction memory, and RE activity modulates mPFC-dHPC synchronization at this frequency [21]. In line with our hypothesis, 8-Hz stimulation of the RE may facilitate memory processes by synchronizing the mPFC-dHPC network throughout the extinction training session.

To examine this, we next performed optogenetic 8-Hz stimulation of the RE in a subset of animals, while recording LFPs in the mPFC and dHPC. Adult male and female rats (*n* = 8, 2 males and 6 females) were injected with AAVs expressing the ChR2 or mCherry (*n* = 4, 1 male and 3 females / group) and implanted with a single fiber optic in the RE. The mPFC and dHPC were implanted with multielectrode arrays comprising sixteen channels covering the PL and IL cortices of the mPFC (8 channels per subregion), and the CA1 region of the dHPC (Fig. S2 and Fig. S3 A-B). Animals first underwent a baseline recording session in a neutral context (Context B) with five 10 s-long 8-Hz stimulation periods. Next day, they were conditioned with five CS-US pairings in Context A. After exposure to the conditioning context the next day, they returned to Context B for a cued extinction session in which the RE was stimulated at 8-Hz during each CS. They were then tested for extinction retrieval the next day, without RE stimulations (Fig. S3 C).

We first examined the effects of stimulation on mPFC and dHPC oscillatory dynamics in a baseline recording session, conducted prior to conditioning. Stimulation was delivered alone, without being paired with CSs. We then examined the effects of stimulation during extinction training, in which the stimulations were paired with CSs. As shown in Figures 2 and 3 A-F (baseline recording and extinction training sessions, respectively), across all successive trials, RE stimulation clearly induced sustained 8-Hz activity in the IL, PL, and dHPC of animals in the ChR2 group compared to animals in the mCherry group. The increased recruitment of the IL, PL, and dHPC was statistically significant relative to the null distribution in ChR2 animals, while mCherry animals did not differ from the null distribution (Figs. 2 and 3 G-I; *p* <.0001. See Source Data Tables). Specifically, during the baseline recording session, the ChR2 group exhibited significantly higher activity with stimulations (Laser on) in the IL, PL, and dHPC compared to the mCherry group (Mann–Whitney *U* test, *p* =.02; Rosenthal’s, *r* =.8165), but did not differ from the mCherry group during the prestimulation baseline and laser off periods (Fig. 2 J-L). Furthermore, the analysis of variance across the extinction session revealed a significant group effect on 8-Hz activity when ranks were aligned on time trials (pre-stimulation baseline, laser on, laser off). This effect was consistent across all analyzed substrates (IL ARS: *F*_1, 62_ = 207.84, *p* <.0001; PL ARS: *F*_1, 62_ = 204.91, *p* <.0001; dHPC ARS: *F*_1, 62_ = 47.77, *p* <.0001) (Fig. 3 J-L). Overall, optogenetic 8-Hz stimulation of the RE induces activity at this frequency in both the mPFC and dHPC.

**Figure 2.**
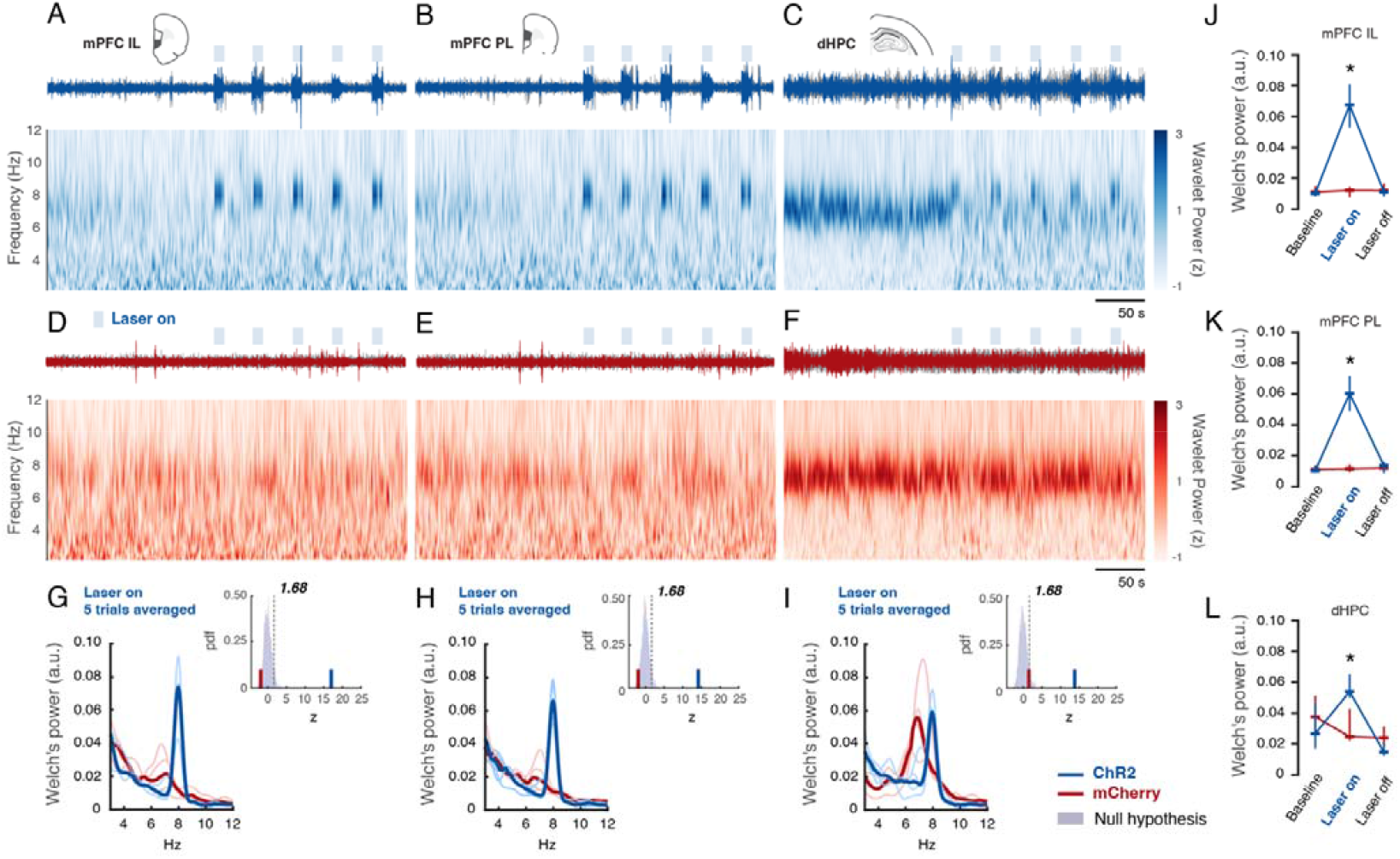
Optogenetic RE stimulation enhances mPFC–dHPC network activity at 8-Hz during the baseline recording session. **(A-F)** Representative filtered signal (7.5 – 8.5 Hz) (IL, PL, and dHPC) from the baseline recording session (upper panels) and time-frequency representation computed using a 1-D wavelet transform (bottom panels), with values expressed as the group median (*n* = 4 / group). Wavelets were *Z*-score normalized across the frequency domain for the ChR2 (in blue) and mCherry (in red) groups, respectively. **(G-I)** Power spectral density with values expressed as the group median (bold lines) and each experimental animal (thin lines) (*n* = 4 / group). Each inserted panel represents the group’s null distribution, and the observed values were transformed into *Z*-scores relative to this distribution. *Z*-values greater than 1.68 were considered statistically significant (one-tailed; *p* < 0.05). **(J-L)** Normalized power around 8 (+/−0.2) Hz across trials (pre-stimulation baseline - laser on - laser off). The ChR2 group exhibited significantly higher activity over stimulations (Laser on) in all recorded substrates (mPFC-IL, mPFC-PL, dHPC) compared to the mCherry group (Mann–Whitney *U* test, *p* =.02; Rosenthal’s, *r* =.8165), and there was no difference during the pre-stimulation baseline and laser off periods. Data are represented as median and 25–75% interquartile range.

**Figure 3.**
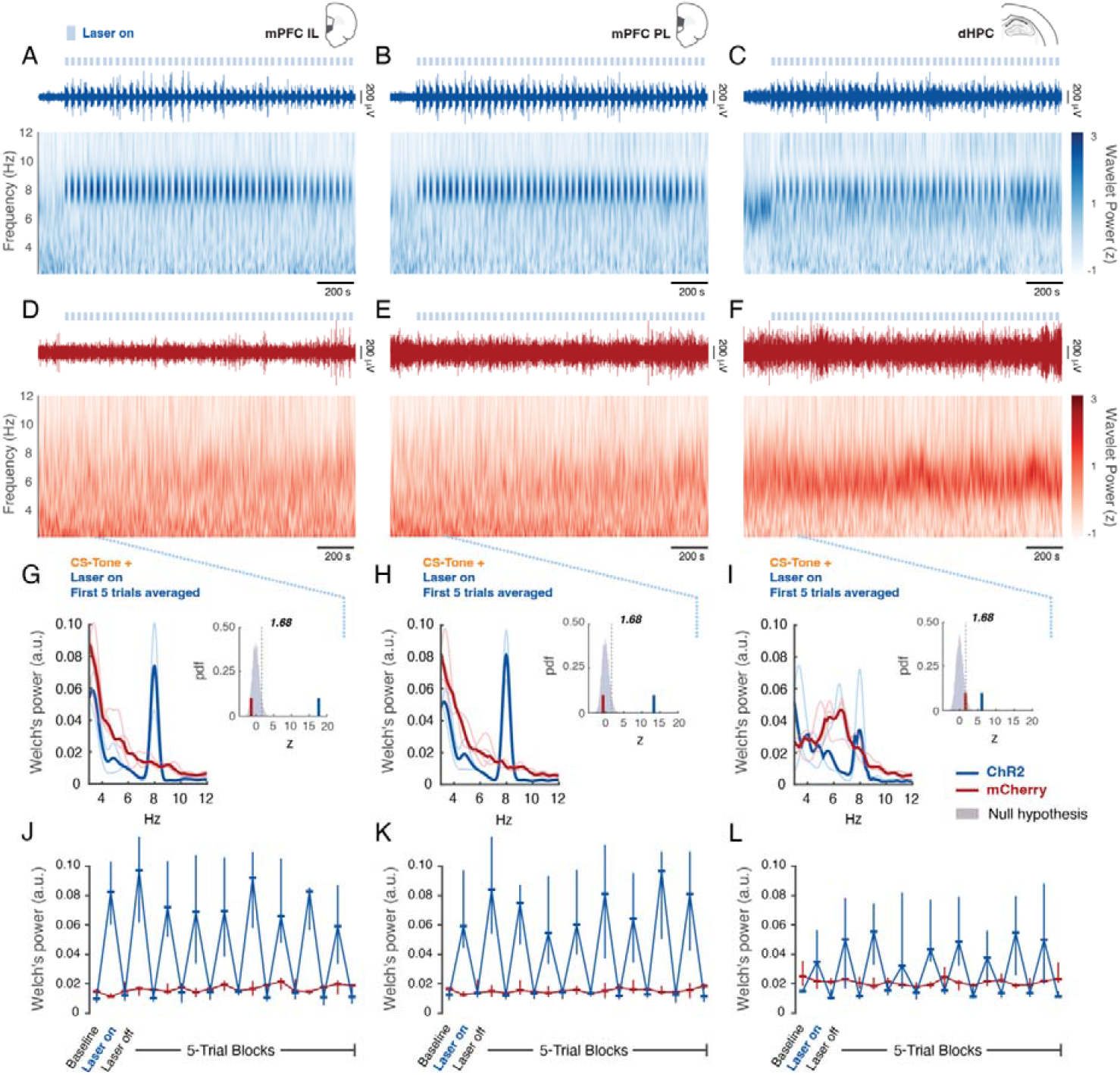
Optogenetic RE stimulation enhances mPFC–dHPC network activity at 8-Hz during the extinction session. **(A-F)** Representative filtered signal (7.5 – 8.5 Hz) from the extinction recording session (upper panels) and time-frequency representation computed using a 1-D wavelet transform (bottom panels), with values expressed as the group median (*n* = 4 / group). Wavelets were *Z*-score normalized across the frequency domain for the ChR2 (in blue) and mCherry (in red) groups, respectively. **(G-I)** Power spectral density with values expressed as the group median (bold lines) and each experimental animal (thin lines) (*n* = 4 / group). Each inserted panel represents the group’s null distribution, and the observed values were transformed into *Z*-scores relative to this distribution. *Z*-values greater than 1.68 were considered statistically significant (one-tailed; *p* < 0.05). **(J-L)** Normalized power around 8 (+/− 0.2) Hz across trials (pre-stimulation baseline - laser on - laser off). There was a significant main effect of group over stimulation periods (Laser on) in all recorded substrates (mPFC-IL, mPFC-PL, dHPC) compared to the mCherry animals (IL ARS: *F*_1, 62_ = 207.84, *p* <.0001; PL ARS: *F*_1, 62_ = 204.91, *p* <.0001; dHPC ARS: *F*_1, 62_ = 47.77, *p* <.0001). Data are represented as median and 25–75% interquartile range.

### Theta-paced stimulation of the RE increases the phase coherence in the mPFC-dHPC network

Because significant activity was observed in the mPFC and dHPC at the stimulation frequency of 8-Hz, an additional analysis was performed to assess the relative phase coherence between the dHPC and both the PL and IL, aiming to characterize their interactions during stimulation at both dynamic and temporal scales [30, 34, 35].

The RE stimulation promoted an enhanced temporal synchrony between dHPC and both the PL and IL regions for the ChR2 group, as measured by phase coherence using phase-locking values (PLV). This pattern of increased synchrony was observed across all stimulation trials in the ChR2 animals during the baseline (Fig. 4 A-D) and extinction (Fig. 5 A–D) recording sessions, in contrast to the mCherry control animals (Figs. 4&5 E–H). Moreover, looking at the first and last five trials of the extinction training, we observed that the temporal synchrony between the IL-dHPC and PL-dHPC was sustained during the whole stimulation period (20 s, laser on) for ChR2 animals (Fig. S4), but not for mCherry animals (Fig. S5). The sustained phase synchrony between the IL-dHPC and PL-dHPC was confirmed by the larger effect relative to the null distribution in ChR2 animals (Figs. 4 and 5 I-J for baseline recording and extinction recording, respectively. Also see Source Data Tables). It is important to emphasize that the mCherry group, too, fell outside of the null distribution. Therefore, a certain level of synchrony between IL-dHPC and PL-dHPC may still be present for mCherry animals, particularly given that the dHPC has monosynaptic projections to the mPFC [36, 37]. However, in a pairwise comparison during the baseline recording session, the ChR2 group exhibited significantly higher PLV values compared to the mCherry group (laser on) (Mann–Whitney *U* test, *p* =.02; Rosenthal’s *r* =.8165). No significant differences were observed between groups during the pre-stimulation baseline or laser off periods (Fig. 4 K–L). Finally, the analysis of variance across the extinction session also revealed a strong group effect on PLV at 8-Hz when data were aligned across trial phases (pre-stimulation baseline, laser on, laser off) with a consistent effect for all comparisons (Fig. 5 K-L, IL-dHPC ARS: *F*_1, 62_ = 41.22, *p* <.0001; PL-dHPC ARS: *F*_1, 28_ = 47.33, *p* <.0001).

**Figure 4.**
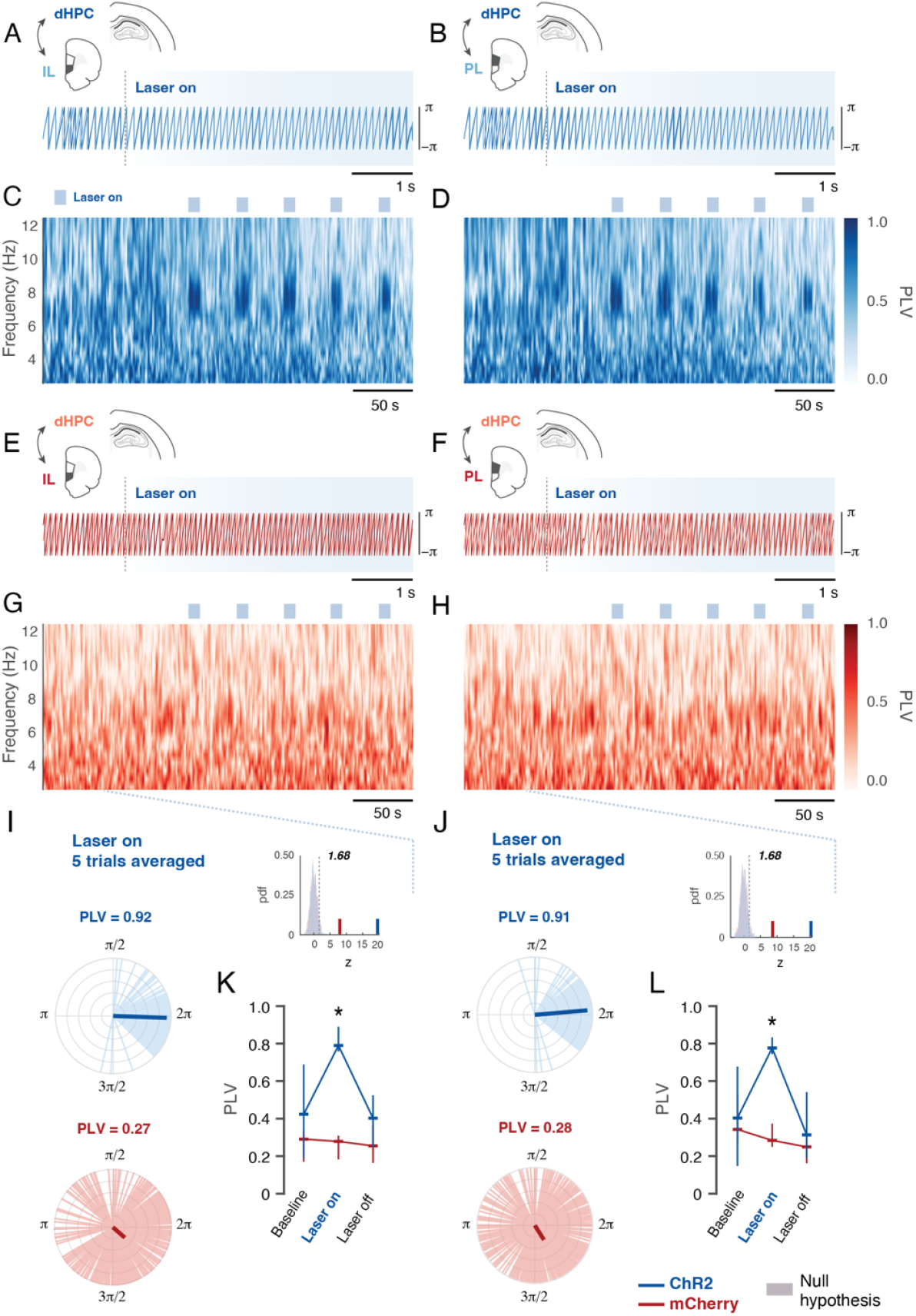
Optogenetic RE stimulation enhances mPFC–dHPC phase coherence at 8-Hz during the baseline recording session. **(A-D)** Representative relative phase signal (7.5 – 8.5 Hz; Laser onset), extracted using the Hilbert transform from the baseline recording session and corresponding phase-locking values (PLV) between 2-12 Hz for IL-dHPC **(A&C)** and PL-dHPC **(B&D)** pairs for ChR2 animals (*n* = 4). **(E-H)** Same for mCherry animals (*n* = 4). A PLV of 0 indicates a random or uniform phase distribution, whereas a PLV of 1 reflects perfect phase synchrony between the recorded substrates. **(I-J)** Polar plots indicating the distribution of relative phase differences for ChR2 group (light blue lines) and mCherry group (light red lines) while the bold lines indicate the median PLV magnitude expressed as the resultant mean phase vector over the 5 trials (*n* = 4 / group). Each inserted panel represents the group’s null distribution, and the observed values were transformed into *Z*-scores relative to this distribution. *Z*-values greater than 1.68 were considered statistically significant (one-tailed; *p* < 0.05). **(K-L)** PLV values across trials (pre-stimulation baseline - laser on - laser off). The ChR2 group exhibited significantly higher synchronism over stimulation period (Laser on) in both IL-dHPC **(K)** and PL-dHPC **(L)** compared to the mCherry group (Mann–Whitney *U* test, *p* =.02; Rosenthal’s, *r* =.8165). There is no difference during the pre-stimulation baseline and laser off periods. Data are represented as median and 25–75% interquartile range.

**Figure 5.**
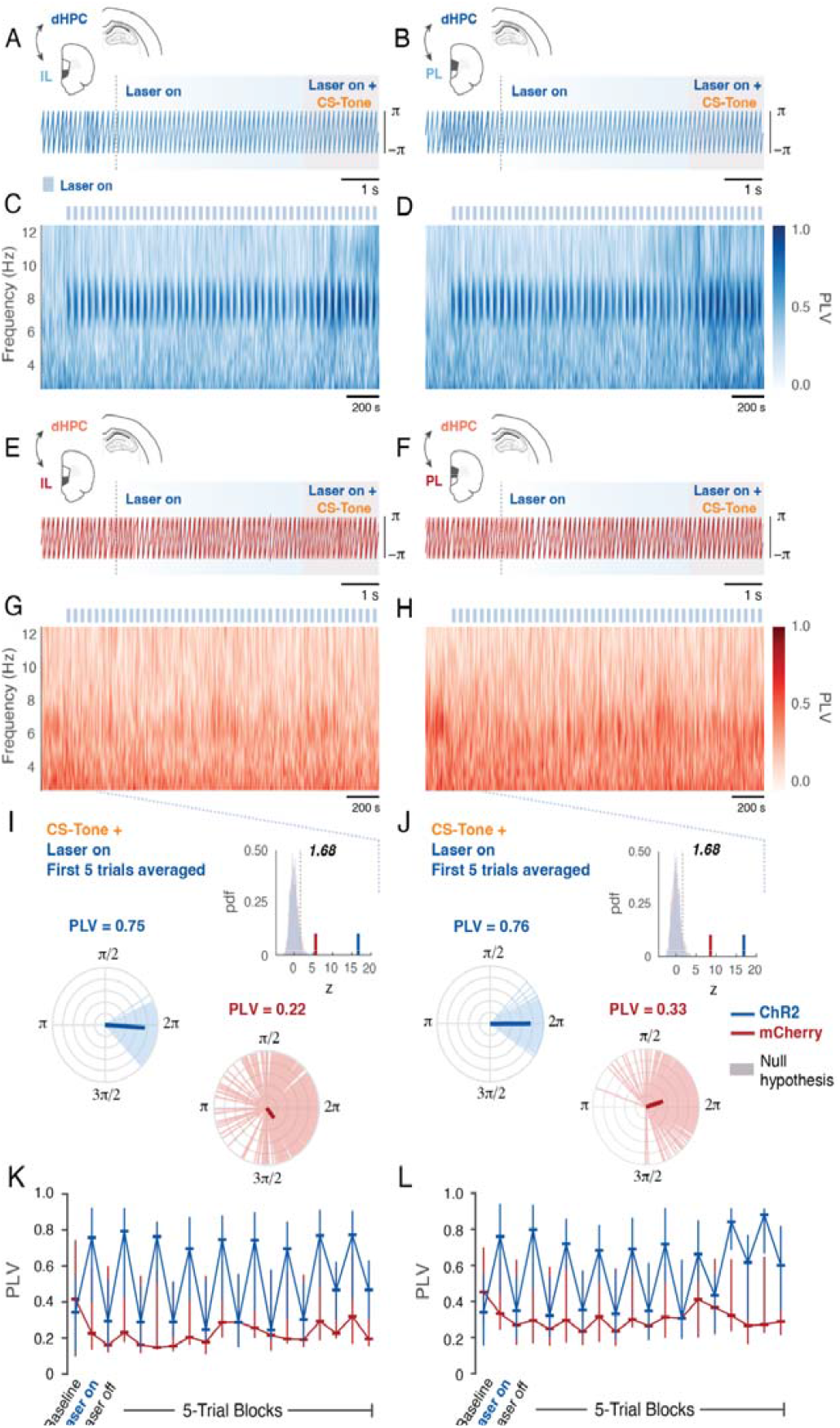
Optogenetic RE stimulation enhances mPFC–dHPC phase coherence at 8-Hz during the extinction session. **(A-H)** Representative relative phase signal (7.5 – 8.5 Hz; Laser onset), extracted using the Hilbert transform from the extinction session and corresponding phase-locking values (PLV) between 2-12 Hz for IL-dHPC **(A&C)** and PL-dHPC **(B&D)** pairs for ChR2 animals (*n* = 4). **(E-H)** Same for mCherry animals (*n* = 4). A PLV of 0 indicates a random or uniform phase distribution, whereas a PLV of 1 reflects perfect phase synchrony between the recorded substrates. **(I-J)** Polar plots indicating the distribution of relative phase differences for ChR2 group (light blue lines) and mCherry group (light red lines) while the bold lines indicate the median PLV magnitude expressed as the resultant mean phase vector over the first 5-trial block (*n* = 4 / group). Each inserted panel represents the group’s null distribution, and the observed values were transformed into *Z*-scores relative to this distribution. *Z*-values greater than 1.68 were considered statistically significant (one-tailed; *p* < 0.05). **(K-L)** PLV values across 5-trial blocks (prestimulation baseline - laser on - laser off). The ChR2 group exhibited a significant main group effect over stimulation period (Laser on) for both IL-dHPC **(K)** and PL-dHPC **(L)** compared to the mCherry group (IL-dHPC ARS: *F*_1, 62_ = 41.22, *p* <.0001; PL-dHPC ARS: *F*_1, 28_ = 47.33, *p* <.0001). Data are represented as median and 25–75% interquartile range.

Collectively, these findings indicate that RE optogenetic stimulation at 8-Hz consistently entrains neural activity at this frequency. Our results suggest a prolonged functional coupling between the dHPC and mPFC with RE-driven activation, during both pre-conditioning baseline and post-conditioning extinction training sessions.

## DISCUSSION

Here, we showed that theta-paced optogenetic stimulation of the RE during extinction training suppresses conditioned freezing and increases grooming behavior in rats. Stimulation of the RE facilitated the acquisition of long-term extinction memories, as evidenced by low freezing levels during a stimulation-free extinction retrieval test. Moreover, we demonstrated that 8-Hz RE stimulation induced 8-Hz activity in both the PL and IL cortices of the mPFC and HPC and synchronized these structures at this stimulation frequency. These results suggest that RE modulation of mPFC-HPC oscillatory dynamics may be important for extinction memory.

Previous research has pointed to a theta oscillatory coupling between the mPFC and HPC during extinction retrieval [21, 38, 39]. Although there is an HPC lead of PL theta oscillations during reexposure to the extinction context in extinction retrieval testing [40], IL leads HPC theta oscillations during extinction retrieval CSs [39]. This is in line with the “retrieval suppression” hypothesis, which proposes PFC control over HPC during memory retrieval. As there are few direct mPFC⍰HPC projections, cortical control over the hippocampal processes is likely modulated by the RE [6, 11–13, 16]. Importantly, Totty and colleagues [21] revealed that pharmacological inactivation of the RE impairs mPFC-HPC theta coherence (6 - 9 Hz) as well as the retrieval of extinction memories. Current results extend on these findings by showing that extinction memory can be facilitated by theta-paced stimulations of the RE, which, in turn, synchronize the mPFC-HPC network. Thus, mPFC-HPC theta oscillatory coherence, mediated by the RE, is key for extinction memory.

It should be noted that our results do not address the source of theta rhythm in the RE. Whether these oscillations emerge from local theta generators or are volume-conducted to the RE from elsewhere remains unclear. The RE forms asymmetrical synapses on both the pyramidal cells and inhibitory interneurons in the mPFC [41, 42] and the CA1 of the HPC [43, 44]. Despite this known connectivity pattern, the network cellular mechanism by which the RE impacts oscillatory dynamics in the mPFC and dHPC is not fully understood. In addition, reports have been contradictory regarding the effect of RE stimulation on HPC neuronal excitability [45, 46] and the role of RE in synchronizing the mPFC-HPC network at theta frequencies. Although some work points to a role for RE in mPFC-HPC theta coherence [21, 47–50], others have failed to show such a role [51, 52]. Ito et al. [53] reported that theta oscillatory coherence in the mPFC-RE-dHPC circuit is essential for information flow and that the supramammillary nucleus of the hypothalamus coordinates this coherence and gates the information flow from the mPFC to dHPC via the RE. This work suggests an external structure controlling temporal coordination in the mPFC-RE-dHPC network. It should be noted that the aforementioned studies employed different tasks, which may account for some of the differences in results. However, more work is clearly needed to characterize the network and oscillatory dynamics between the RE, mPFC, and HPC.

Recently, Goh et al. [27] reported that optogenetic stimulation of the RE causes compulsive-like grooming behavior in rats. We confirmed this finding in a subset of ChR2 animals. However, despite the small number of ChR2 animals observed, our results suggest that grooming behavior caused by RE photostimulation does not explain the effects of photostimulation on extinction learning or extinction retrieval, as the animal showing consistent photostimulation-induced grooming throughout extinction showed poorer extinction retrieval compared to the other, which stopped grooming in the later trials of extinction. Moreover, grooming behavior is not associated with strong hippocampal theta oscillations [54, 55]. Therefore, animals’ grooming while showing strong 8-Hz power in the mPFC and HPC throughout the stimulation period is unexpected. This may suggest parallel mechanisms and different circuits engaged with RE theta-paced stimulations. Indeed, Goh et al. [27] revealed that RE to dorsal premammillary nucleus projections mediate grooming behavior. Because the present study involved stimulation of the RE as a whole, we are likely capturing both dorsal premammillary nucleus-projecting neurons of the RE as well as the mPFC- and HPC-projecting RE neurons and those that receive projections from the mPFC and HPC. Different populations of RE neurons may give rise to different behavioral outputs depending on their connectivity with other regions.

Goh et al. [27] also reported increased avoidance behavior and anxiety with RE stimulation. In a place preference task, rats avoided the compartment paired with RE photo-stimulation. In an open field test, they spent less time in the center and showed increased freezing following RE stimulation. In addition, in an elevated plus maze, they spent less time exploring the open arms following RE stimulation [27]. These findings led Goh et al. [27] to conclude that RE photo-stimulation is aversive and distressing for the animals, creating anxiogenic effects. However, our results point to a different interpretation. RE theta-paced stimulation suppressed conditioned freezing behavior rather than increasing it, which carried over to the next day in the stimulation-free retrieval test. Although avoidance or anxiety behavior was not explicitly tested in the current study, when animals were returned to the extinction context for retrieval testing, they exhibited very low freezing levels and normal exploratory behavior (see Fig. 1 D, Extinction Retrieval BL freezing level). Our results suggest that RE theta-paced photo-stimulation promotes a low-fear state and facilitates extinction retrieval, rather than driving anxiety and avoidance. Theta-oscillatory coupling between the mPFC and HPC is a correlate of extinction retrieval and low fear states [21, 38–40]. On the other hand, HPC-amygdala theta oscillatory coupling has been shown to emerge during the retrieval of fear memories, promoting a high fear state [56, 57]. However, recognition of safety entrains amygdala neuronal firing to mPFC theta oscillations [58], suggesting that mPFC input to both the HPC and amygdala can be critical for suppressing conditioned fear responses. Karalis et al. [59] demonstrated that the mPFC and amygdala show synchronized 4-Hz oscillations, with mPFC leading the amygdala, which predict fearful freezing behavior. These oscillations are internally generated and distinct from hippocampal theta oscillations, as blocking hippocampal theta does not abolish mPFC 4-Hz oscillations. Moreover, optogenetic stimulation of the mPFC parvalbumin-expressing interneurons induces synchronized 4-Hz oscillations in the mPFC-amygdala network and freezing behavior [59]. Ozawa et al. [33] showed that optogenetically-induced 4-Hz and 8-Hz oscillations in the amygdala increase and decrease freezing behavior, respectively, following extinction training. They reported that in both the mPFC and amygdala, 4-Hz oscillations signal a fearful state, whereas 8-Hz oscillations signal a safety state, and the balance between these oscillations predicts freezing. Totty et al. [21] revealed increased and coherent 8-Hz power in the mPFC and HPC during extinction retrieval. The RE, too, showed increased 8-Hz power in later trials of extinction training compared to earlier trials, when the fear level is lower. Combined, these findings suggest that 8-Hz power in the mPFC, amygdala, and HPC as well as network coherence at this frequency is associated with a low fear state. Our results extend on these findings by showing that mPFC-dHPC 8-Hz coherence induced by optogenetic RE stimulations also promotes a low fear state.

Here, we showed that theta-paced stimulation of the RE during extinction training facilitates the retrieval of extinction memories the next day. One might argue that such stimulation during extinction helps consolidate the extinction memory, leading to stronger or longer-lasting extinction memory. Indeed, there is evidence that the RE is involved in memory consolidation by facilitating mPFC-HPC interaction during sleep [60–62]. Offline replay of newly acquired memories is key for memory consolidation, and it is known to involve a slow oscillatory activity coupling between the mPFC and HPC [63–65]. The RE facilitates information exchange between the prefrontal-hippocampal networks during slow-wave sleep. Inactivating the RE impairs mPFC-HPC oscillatory synchrony during slow oscillations in anesthetized rats [60–62].

Notably, a recent study by Lee et al. [66] demonstrated that optogenetic excitation of RE neurons in mice leads to deep NREM sleep. Moreover, they reported that, prior to sleep, animals exhibited increased grooming, a sleep-preparatory behavior. Authors described an RE-to-zona incerta pathway that undergoes plasticity with sleep deprivation and subsequent homeostatic sleep [66]. These results suggest that the RE has a direct role in sleep, which may further contribute to memory consolidation. However, it should be noted that some studies have failed to find a role for RE in memory consolidation. Mei et al. [67] showed that the RE is not involved in consolidation of spatial memories, though it is critical for the retrieval of spatial memories. Vasudevan et al. [68] revealed that the RE is not involved in either consolidation or reconsolidation of fear extinction memories, whereas Troyner and Bertoglio [69] reported that reconsolidation of contextual fear memories depends on the RE. Although task differences in these studies should be noted, it emerges that more studies are needed to determine the role of the RE in extinction memory consolidation.

Overall, our results suggest that the RE may be a novel therapeutic target for facilitating extinction memories. Theta-paced stimulation of the RE, which enhances long-range oscillations and synchronizes the mPFC-HPC network, may become an important therapeutic tool for strengthening extinction memory. Indeed, RE stimulation could be used as an adjunct or as an alternative to exposure-based therapies to treat people with PTSD and other fear and anxiety disorders. Future efforts should be made to test the effects of RE theta-paced stimulations in humans.

## ACKNOWLEDGEMENTS

This work was supported by the National Institutes of Health (R01MH065961 and R01MH117852 to S.M.).

## AUTHOR CONTRIBUTIONS

TT and SM designed the optogenetic experiment; TT performed the optogenetic experiment and analyzed behavioral data; TT, FM, and SM designed the electrophysiology experiment; TT and FM performed the electrophysiology experiment. FM analyzed the electrophysiology data; TT, FM, and SM wrote the manuscript.

## DATA AVAILABILITY

The data from these experiments are available from the corresponding author upon request.

## SUPPLEMENTARY RESULTS

**Table S1.** Baseline recording grooming (%)

| Animal | Group | BL | S1 | ISI1 | S2 | ISI2 | S3 | ISI3 | S4 | ISI4 | S5 | ISI5 |
| --- | --- | --- | --- | --- | --- | --- | --- | --- | --- | --- | --- | --- |
|  |  | (180s) | (10s) | (30s) |  |  |  |  |  |  |  |  |
| 1 | mCherry | 14.2 | 0 | 0 | 0 | 0 | 0 | 0 | 0 | 0 | 0 | 0 |
| 7 | mCherry | 17.8 | 0 | 0 | 0 | 0 | 0 | 0 | 0 | 0 | 0 | 0 |
| 8 | mCherry | 11.3 | 16.7 | 82.1 | 0 | 0 | 0 | 0 | 0 | 0 | 0 | 0 |
| 9 | mCherry | 17.8 | 0 | 0 | 5 | 47.1 | 17.7 | 0 | 0 | 0 | 0 | 0 |
| 10 | ChR2 | 38.6 | 81.7 | 18.8 | 68.7 | 15.8 | 80.7 | 8.4 | 88.7 | 20.8 | 60.7 | 31.1 |
| 11 | ChR2 | 32.4 | 85.7 | 15.4 | 85.7 | 40.1 | 86.7 | 6.4 | 34 | 0 | 62.7 | 21.1 |

**Table S2.** Extinction recording grooming (%) during the first five trials.

| Animal | Group | BL | S1 | ISI1 | S2 | ISI2 | S3 | ISI3 | S4 | ISI4 | S5 | ISI5 |
| --- | --- | --- | --- | --- | --- | --- | --- | --- | --- | --- | --- | --- |
|  |  | (175s) | (20s) | (20s) |  |  |  |  |  |  |  |  |
| 1 | mCherry | 13 | 0 | 0 | 0 | 0 | 0 | 0 | 0 | 0 | 0 | 0 |
| 7 | mCherry | 13.4 | 0 | 0 | 0 | 0 | 0 | 0 | 0 | 0 | 0 | 0 |
| 8 | mCherry | 35.6 | 0 | 0 | 0 | 0 | 0 | 0 | 0 | 0 | 0 | 0 |
| 9 | mCherry | 24.9 | 0 | 0 | 0 | 0 | 0 | 0 | 0 | 0 | 0 | 0 |
| 10 | ChR2 | 27.6 | 84.8 | 33.5 | 89 | 64.3 | 89 | 100 | 100 | 21.8 | 73.2 | 23.5 |
| 11 | ChR2 | 43.3 | 84.2 | 13.3 | 88.3 | 54.2 | 90 | 78.3 | 84.2 | 85 | 5.9 | 0 |

**Table S3.** Extinction recording grooming (%) during the last five trials.

| Animal | Group | S41 | ISI41 | S42 | ISI42 | S43 | ISI43 | S44 | ISI44 | S45 | ISI45 |
| --- | --- | --- | --- | --- | --- | --- | --- | --- | --- | --- | --- |
|  |  | (20s) | (20s) |  |  |  |  |  |  |  |  |
| 1 | mCherry | 0 | 0 | 0 | 0 | 0 | 0 | 0 | 0 | 0 | 0 |
| 7 | mCherry | 0 | 3.3 | 0 | 0 | 0 | 0 | 0 | 0 | 0 | 0 |
| 8 | mCherry | 26.7 | 0 | 59.8 | 53.5 | 0 | 0 | 0 | 0 | 0 | 0 |
| 9 | mCherry | 0 | 0 | 0 | 0 | 5.8 | 0 | 0 | 0 | 0 | 0 |
| 10 | ChR2 | 0 | 0 | 18.8 | 30.3 | 0 | 0 | 0 | 0 | 0 | 0 |
| 11 | ChR2 | 0 | 87.5 | 100 | 49.8 | 63.5 | 100 | 100 | 57.3 | 64.3 | 17.3 |

**Figure S1.**
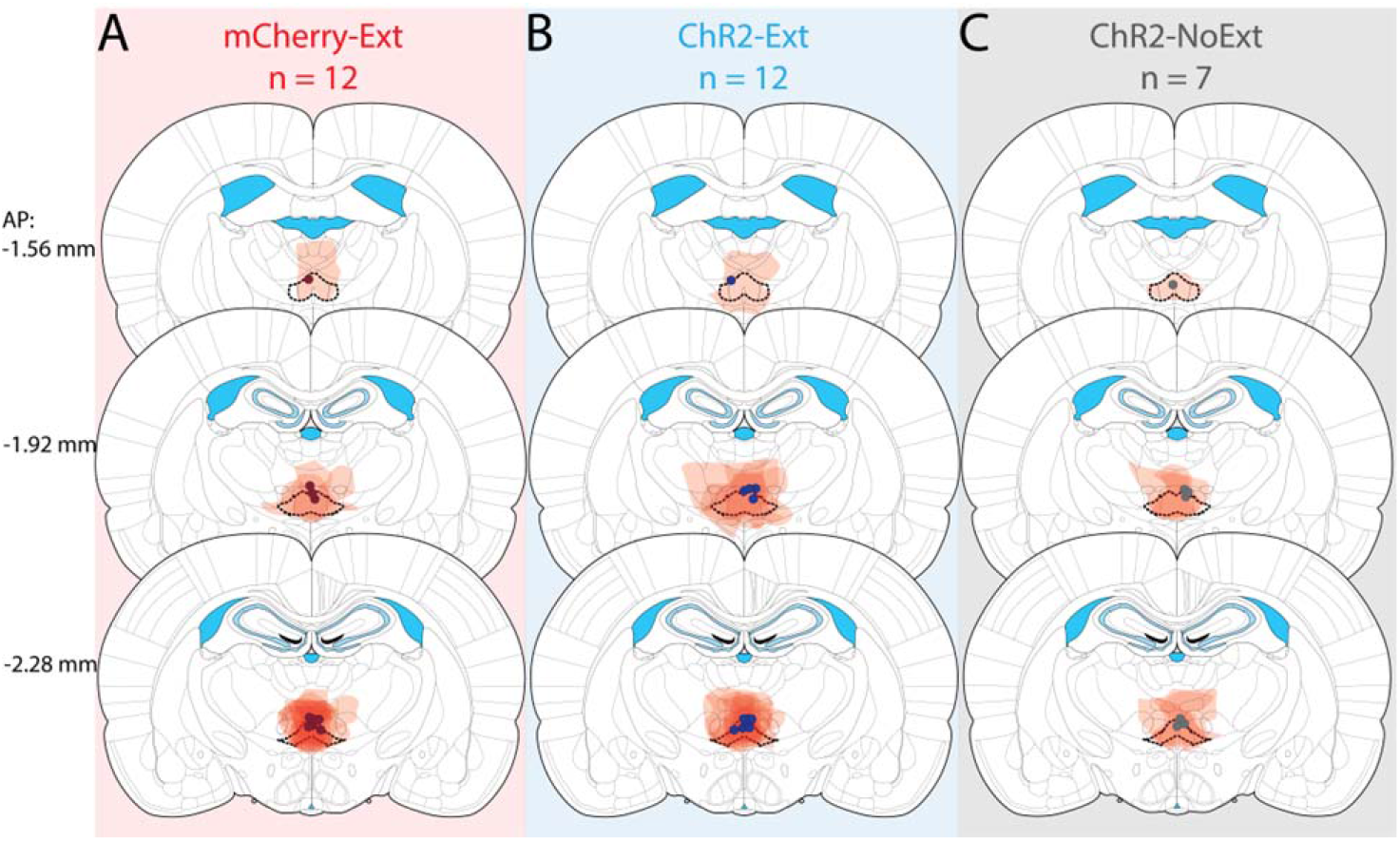
Viral expression and fiber optic optic placements in the RE. Animals were injected either with AAV9-CaMKIIa-ChR2-mCherry (*n* = 19) or AAV9-CaMKIIa-mCherry (*n* = 12) and were implanted with a single fiber optic in the RE. Viral expression and fiber optic placements were verified for mCherry-Ext **(A)**, ChR2-Ext **(B)** and ChR2-NoExt **(C)** groups at three different anterior-posterior (AP) coordinates: −1.56, −1.92, and −2.28 mm. All fiber optic placements (colored dots) were confined to the RE borders which are denoted by thick dashed lines. Stereotaxic atlas positions are adapted from Paxinos and Watson (2014).

**Figure S2.**
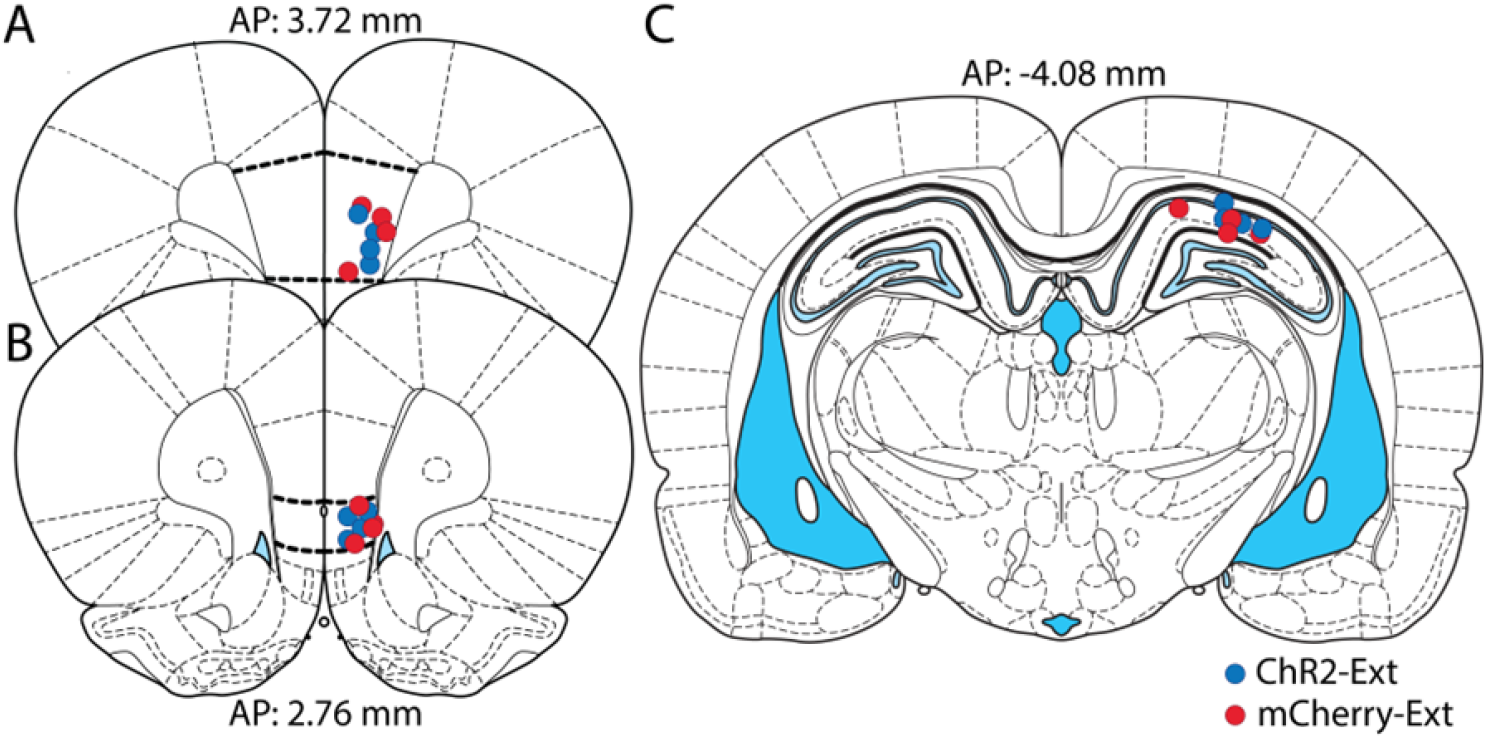
Electrode placements in the mPFC and dHPC. Multi-electrode array placements in the PL **(A)** and IL **(B)** cortices of the mPFC and dorsal CA1 region **(C)** of the HPC in ChR2-Ext (*n* = 4, blue circles) and mCherry-Ext (*n* = 4, red circles) group animals. Stereotaxic atlas positions are adapted from Paxinos and Watson (2014). PL: Prelimbic cortex, IL: Infralimbic cortex, mPFC: Medial prefrontal cortex, and HPC: Hippocampus.

**Figure S3.**
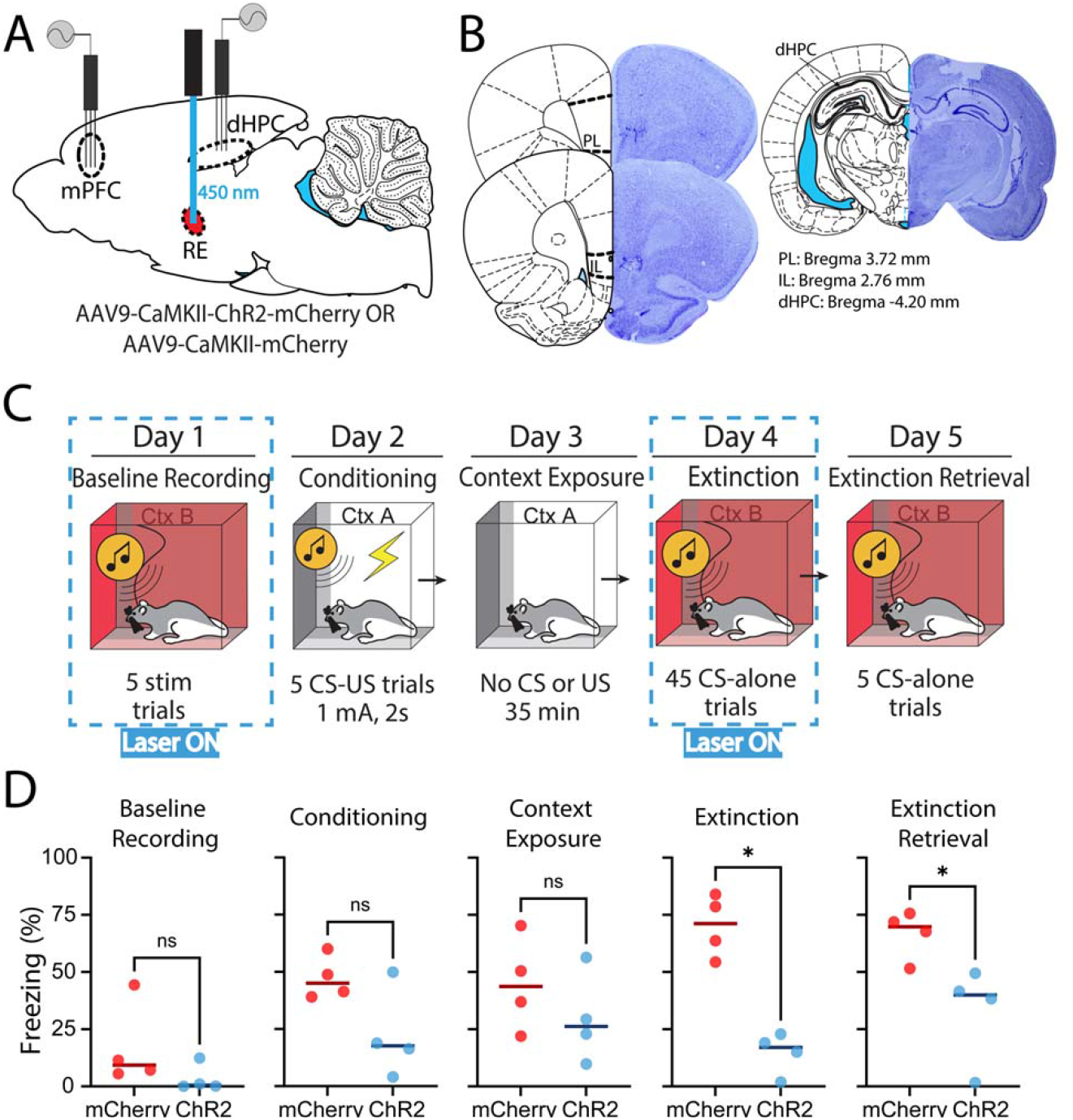
Electrophysiological recording experiment design and behavioral results. **(A)** Surgical procedure. Animals were injected either with AAV9-CaMKIIa-ChR2-mCherry or AAV9-CaMKIIa-mCherry (*n* = 4 / group, 1 male and 3 females) and implanted with a single fiber optic in the RE. mPFC and dHPC were implanted with multielectrode arrays. **(B)** Representative electrode placement from the IL and PL cortices of the mPFC and the dHPC. **(C)** Behavioral protocol used. The same behavioral protocol was used as in the optogenetics experiment, but animals additionally underwent a baseline recording session with five 10 s-long 8-Hz stimulations of the RE separated by 30 s ISIs. **(D)** Behavioral data from the mCherry and ChR2 group animals in each day. mCherry and ChR2 animals showed comparable freezing levels during baseline recording, conditioning, and conditioning context exposure (Mann-Whitney test, *p*s >.05). ChR2 animals demonstrated significantly lower freezing levels compared to mCherry animals during both extinction and extinction retrieval (Mann-Whitney test, *p*s =.0286). Lines indicate median value in scatter plots in **(D)**. Stereotaxic atlas positions are adapted from Paxinos and Watson (2014). PL: Prelimbic cortex, IL: Infralimbic cortex, mPFC: Medial prefrontal cortex, and dHPC: Dorsal hippocampus.

**Figure S4.**
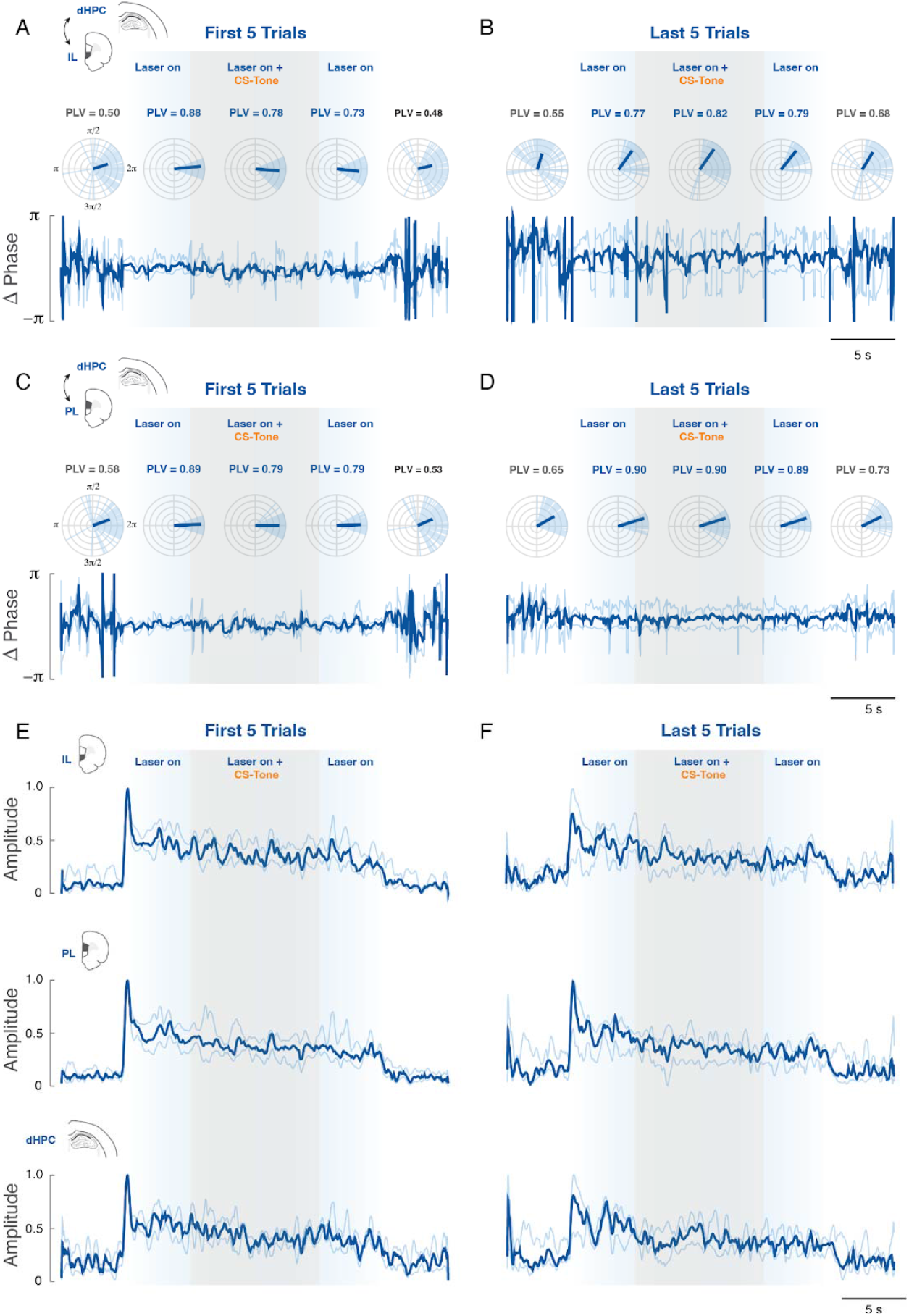
RE stimulation induces sustained 8-Hz activity in the ChR2 group. During the extinction session, animals were exposed to a series of CS presentations, with 8-Hz optogenetic stimulation delivered during the CS periods. The analysis window included: 5 s before optogenetic stimulation onset (Laser off), 5 s of optogenetic stimulation alone prior to CS onset (Laser on), 10 s of CS presentation with concurrent optogenetic stimulation (Laser on + CS trials), the final 5 s of optogenetic stimulation alone after CS offset (Laser on), and five seconds following optogenetic stimulation offset (Laser off). **(A**,**C**,**E)** Results from the first five trials of extinction. **(B**,**D**,**F)** Results from the last five trials of extinction. **(A-D)** Polar plots (upper panels) indicate the distribution of relative phase differences for ChR2 animals (light blue lines) while bold lines indicate the median PLV magnitude expressed as the resultant mean phase vector over the 5-trial block (*n* = 4). Relative phase differences (7.5 – 8.5 Hz) over time expressed as median and 25–75% interquartile range. **(E-F)** Amplitude envelope (7.5 – 8.5 Hz) for IL, PL, and dHPC expressed as median and 25–75% interquartile range, respectively.

**Figure S5.**
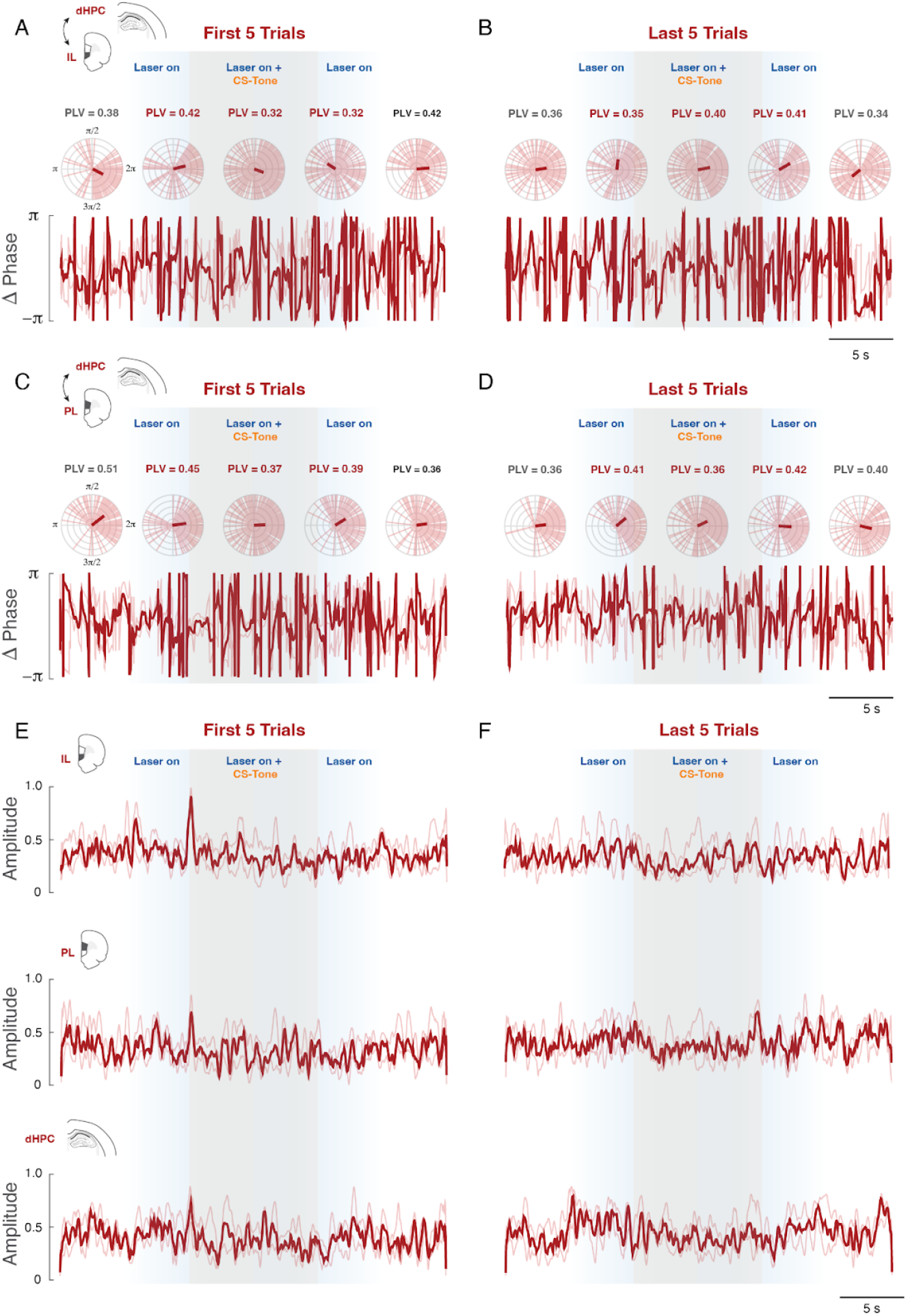
RE stimulation does not affect the mPFC-dHC network activation and synchronization in the mCherry group. The analyses explained in SupplementaryFigure 5 were repeated in the mCherry control group. The sustained activity observed in the ChR2 group was absent in the mCherry group, suggesting that the effects of RE optogenetic stimulation was not non-specific. **(A-D)** Polar plots (upper panels) indicate the distribution of relative phase differences for mCherry animals (light red lines) while the bold lines indicate the median PLV magnitude expressed as the resultant mean phase vector over the 5-trial block (*n* = 4). Relative phase differences (7.5 – 8.5 Hz) over time expressed as median and 25–75% interquartile range. **(E-F)** Amplitude envelope (7.5 – 8.5 Hz) for IL, PL, and dHPC expressed as median and 25–75% interquartile range, respectively.

